# Palmitoylation regulates human serotonin transporter activity, trafficking, and expression and is modulated by escitalopram

**DOI:** 10.1101/2023.05.09.540092

**Authors:** Christopher R. Brown, James D. Foster

**Author notes:** To whom correspondence should be addressed: James D. Foster, Ph.D., Department of Biomedical Sciences, University of North Dakota School of Medicine and Health Sciences, Grand Forks, ND 58202-9037.

## Abstract

In the central nervous system, serotonergic signaling modulates sleep, mood, and cognitive control. During neuronal transmission, the synaptic concentration of serotonin is tightly controlled in a spatial and temporal manner by the serotonin transporter (SERT). Dysregulation of serotonergic signaling is implicated in the pathogenesis of major-depressive, obsessive-compulsive, and autism-spectrum disorders, which makes SERT a primary target for prescription therapeutics, most notably selective-serotonin reuptake inhibitors (SSRIs). S-palmitoylation is an increasingly recognized dynamic post-translational modification, regulating protein kinetics, trafficking, and localization patterns upon physiologic/cellular stimuli. In this study, we reveal that human SERTs are a target for palmitoylation, and using the irreversible palmitoyl acyl-transferase inhibitor, 2-bromopalmitate (2BP) we have identified several associated functions. Using a lower dose of 2BP in shorter time frames, inhibition of palmitoylation was associated with reductions in SERT V_max_, without changes in K_m_ or surface expression. With higher doses of 2BP for longer time intervals, inhibition of palmitoylation was consistent with the loss of cell surface and total SERT protein, suggesting palmitoylation is an important mechanism in regulating SERT trafficking and maintenance of SERT protein through biogenic or anti-degradative processes. Additionally, we have identified that treatment with the SSRI escitalopram decreases SERT palmitoylation analogous to 2BP, reducing SERT surface expression and transport capacity. Ultimately, these results reveal palmitoylation is a major regulatory mechanism for SERT kinetics and trafficking and may be the mechanism responsible for escitalopram-induced internalization and loss of total SERT protein.

## Introduction

Serotonin (5-hydroxytryptamine; 5HT) is a monoamine neurotransmitter responsible for diverse physiologic consequences. Peripherally these include modulation of primary hemostasis, gastrointestinal motility, and fetal organ maturation. In the central nervous system (CNS), 5HT influences sleep, mood, and cognitive control, with dysregulation promoting the development of major-depressive, obsessive-compulsive, and autism-spectrum disorders. CNS serotonergic pathways begin with the Raphe Nuclei, projecting long axons in ascending and descending patterns that diffuse throughout the cerebral cortex and spinal cord. Depolarization of serotonergic neurons initiates the process of synaptic 5HT release, where it can diffuse and activate 5HT receptors on presynaptic and postsynaptic neurons. The serotonin transporter (SERT) is a presynaptic transmembrane protein that functions as the primary mechanism for termination of 5HT signaling, actively removing 5HT from the synaptic cleft into the presynaptic neuron. SERT is dynamically regulated by numerous post-translational modifications that control its trafficking, activity and expression profiles (1-5). Dysregulated control of SERT trafficking and activity has been demonstrated to be a central mechanism in the pathogenesis of autism and obsessive-compulsive disorders (6-9). In this regard, SERT is a primary target of interest in the development of therapeutics to combat the diversity of psychiatric disorders associated with abnormal serotonergic signaling.

The most common class of prescription anti-depressants are selective serotonin reuptake inhibitors (SSRIs). SSRIs function to inhibit SERT by directly blocking the reuptake of 5HT from the synapse, transiently raising the extracellular concentration of 5HT (10). This process results in immediate elevations of synaptic 5HT; however, SSRIs have been reported to require at least one week to three months of constant dosing to achieve clinically observable neuro-cognitive effects, suggesting that the therapeutic potential for SSRIs require longer-term adaptations to the serotonergic system (11). Currently, the therapeutic mechanism of action for this class of antidepressants has yet to be fully uncovered (12,13). *In vitro*, others have reported that the density of cell surface and total cellular SERT is decreased with long-term escitalopram treatments (14,15). When treated with 500 nM escitalopram for three hours, SERT was internalized from the cell surface by 40% in serotonergic neurons (14). This process was found to be reversible when the drug-treated medium was washed away and re-incubated with treatment-free medium (14). Notably, this was performed in the presence of cycloheximide, an inhibitor of protein synthesis, suggesting that the enhanced presence of cell surface SERT originated from an existing intracellular pool of transporter, rather than newly synthesized SERT. This unique finding suggested to us that a reversible post-translational mechanism may be responsible for the observed alterations in SERT trafficking mediated by escitalopram.

However, 12 hours of continuous treatment with 500 nM escitalopram drove nearly all of surface localized SERT to be internalized (14). Additionally, rats treated for 7 days with paroxetine or sertraline had 80-90% reduced total SERT density in Raphe serotonergic neurons (15). Notably, treatment with the monoamine oxidase inhibitor (MAO) phenelzine or the tricyclic antidepressant (TCA) desipramine did not reduce the total cellular density of SERT, but in some cases increased it (15). Another rat-based study revealed that 21 days of paroxetine-treatment followed with a 48-h washout induced a 50-60% reduction in hippocampal and dorsal raphe [^3^H]5HT uptake (16). In addition, the same study determined that when treated with [^3^H]paroxetine, SERT K_d_ values were unchanged, but B_max_ values were reduced by 60-70% (16). Collectively, these data suggest that SSRIs function in the long-term to downregulate SERT *via* mechanisms that coexist with acute blockade of 5HT uptake.

Similar to the process of phosphorylation, S-palmitoylation is a dynamic and reversible post-translational modification that controls protein kinetics, trafficking, and localization in response to current physiologic stimuli (17,18). Palmitoylation has been identified as essential in proper biogenesis of the cystic fibrosis transmembrane regulator (19) and normal trafficking of the immune system toll-like receptor (TLR) pathway regulatory proteins TLR2, 5, 10, MyD88 and Lyn (20), emphasizing how important this modification is in maintaining normal cellular and physiologic function. Previously, we have reported that the dopamine transporter (DAT) is a target for palmitoylation (21). We have identified that palmitoylation of DAT regulates a diverse range of functions including DAT transport kinetics, degradation, and protein kinase C-dependent regulation (21). In this study, we expanded our scope to include SERT and its regulation. Our analysis revealed that SERT is palmitoylated and is sensitive to inhibition of palmitoylation with the irreversible palmitoyl acyltransferase (DHHC) inhibitor, 2-bromopalmitate (2BP). In acute time intervals with lower concentrations, we observed losses of SERT palmitoylation *via* 2BP that were consistent with reductions in SERT V_max_, without changes in SERT K_m_ or cell surface expression. In longer time intervals with higher concentrations, we observed that inhibition of SERT palmitoylation was accompanied with losses in total SERT expression, suggesting that palmitoylation regulates the biogenesis and/or degradation of SERT protein. Importantly, we show that a commonly prescribed antidepressant, escitalopram, inhibits SERT palmitoylation and decreases its surface expression and transport capacity.

## Results

### Identification of SERT Palmitoylation and Inhibition by 2BP

In our initial experiments, we used LLC-PK_1_ or HEK293 cells stably expressing wildtype (WT) HA-hSERT or hSERT, respectively. We utilized LLC-PK_1_ as our primary model system due to our previous success with this cell line in the characterization of DAT palmitoylation (21,23). We subjected membranes prepared from these cells to assess SERT palmitoylation using ABE followed by SDS-PAGE and immunoblotting for SERT. These data revealed hSERT is palmitoylated in both cell lines using anti-HA or anti-hSERT (ST51-2) antibodies for detection (*Fig. 1A*). In the ABE method, hydroxylamine (NH_2_OH) chemically removes palmitate from palmitoylated proteins, allowing for biotinylation with a sulfhydryl specific biotin reagent followed by pull-down with Neutravidin resin and detection by immunoblotting (*Fig. 1A, upper panels - NH_2_OH lanes*). Samples treated with Tris buffer in parallel (*Fig. 1A, upper panels - Tris lanes*) serve as a negative control where palmitate is not removed and is not replaced with biotin, resulting in negligible nonspecific SERT detection by immunoblotting. The lower panel in Figure 1A (*IB*) represents the total SERT level of each sample in aliquots taken from the sample just prior to pull-down with Neutravidin resin. These levels are used to normalize quantification of palmitoylation in the upper panels (*ABE*) where palmitoylated SERT is a fraction of the total SERT present in the sample.

**FIGURE 1.**
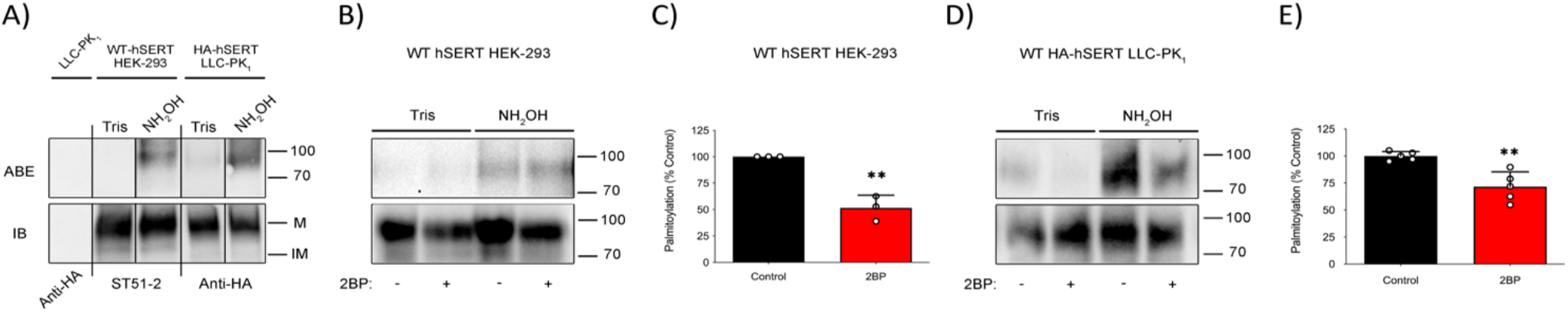
S-palmitoylation of SERT and inhibition by 2BP. *A,* WT-hSERT HEK-293 cells and HA-hSERT-LLC-PK_1_ cells underwent ABE followed by SDS-PAGE and immunoblotting to determine SERT palmitoylation levels (ABE, top panels) and total SERT levels (bottom panels). Extended black lines (|) on the top and bottom of panel A indicate two separate blots/experiments while black lines within the boundaries of the image indicate the removal of duplicate lanes or rearrangement of lane images from the same blot. M_r_ markers for all blots are shown at right. The validated primary monoclonal antibodies used for immunoblotting were anti-human serotonin transporter (ST51-2) for HEK-293 cells expressing WT-hSERT, and anti-hemagglutinin (Anti-HA) for HA-hSERT detection. WT hSERT HEK-293 cells (B) or WT HA-hSERT LLC-PK_1_ cells (D) underwent ABE after treatment with vehicle or 10 µM 2BP for 18 hours and palmitoylation levels were determined after SDS-PAGE by immunoblotting. *C,* Quantification of hSERT palmitoylation in C (mean ± SD of 3 independent experiments relative to control normalized to 100%.; ** p < 0.01 versus control; Student’s t-test for independent samples). *E,* Quantification of palmitoylation in D (mean ± SD of 5 independent experiments relative to control normalized to 100%; ** p < 0.01 versus control; Student’s t-test for independent samples).

To further characterize SERT palmitoylation, we used an irreversible inhibitor of palmitoyl acyltransferases, 2-bromopalmitate (2BP), to inhibit cellular palmitoylation. We have previously demonstrated with DAT (21) the irreversible nature of 2BP and its impact on DAT activity and trafficking in acute and chronic conditions. In our current study, we treated cells expressing either WT hSERT or HA-hSERT with 10 µM 2BP for 18 h (Fig. 1, B and D respectively). In HEK293 cells expressing WT hSERT, 2BP decreased hSERT palmitoylation to 51.5 ± 11.9% of vehicle treated control cells (Fig. 1B and 1C) (p<0.001 via Student’s t-test, n=3). In LLC-PK_1_ cells expressing HA-hSERT, we found similar results under identical conditions, with HA-hSERT palmitoylation decreased to 71.6 ± 13.6% relative to control (Fig. 1D and 1E) (p<0.01 via Student’s t-test, n=5). These findings demonstrate that 2BP inhibits SERT palmitoylation in two independent heterologous cell lines, revealing that SERT palmitoylation can be inhibited by 2BP similarly to DAT (21).

### Acute Inhibition of SERT Palmitoylation Downregulates Transport Kinetics Without Altering Surface Expression

After we demonstrated that treatment with 2BP can inhibit palmitoylation of SERT, we directed our efforts to determine if SERT palmitoylation was sensitive to acute inhibition (0-120 min) and if this impacted SERT trafficking and/or activity. Our previous studies with DAT demonstrated that treatment with 10 µM 2BP inhibited palmitoylation within 30 min that resulted in reduced DAT V_max_ without changes in surface or total DAT expression (21).

To test this, we treated cells expressing SERT in a time-course fashion with vehicle or 7.5 µM 2BP in 30-min increments for a total of 120 min. ABE analysis of these samples revealed that 30 min of 2BP inhibition reduced SERT palmitoylation to 76.9 ± 15% that continued in a time-dependent fashion through 120 min (50.4 ± 5.3%) (Fig. 2A, p<0.05 and p<0.001 via ANOVA with Tukey post-test, n=3-4, respectively). The time-dependent nature of these results is consistent with the irreversible mechanism of 2BP action for inhibiting palmitoylation (p<0.05 via ANOVA with Tukey post-test). Immunoblotting for total HA-hSERT protein revealed no detectable changes in total SERT protein expression under these conditions.

**FIGURE 2.**
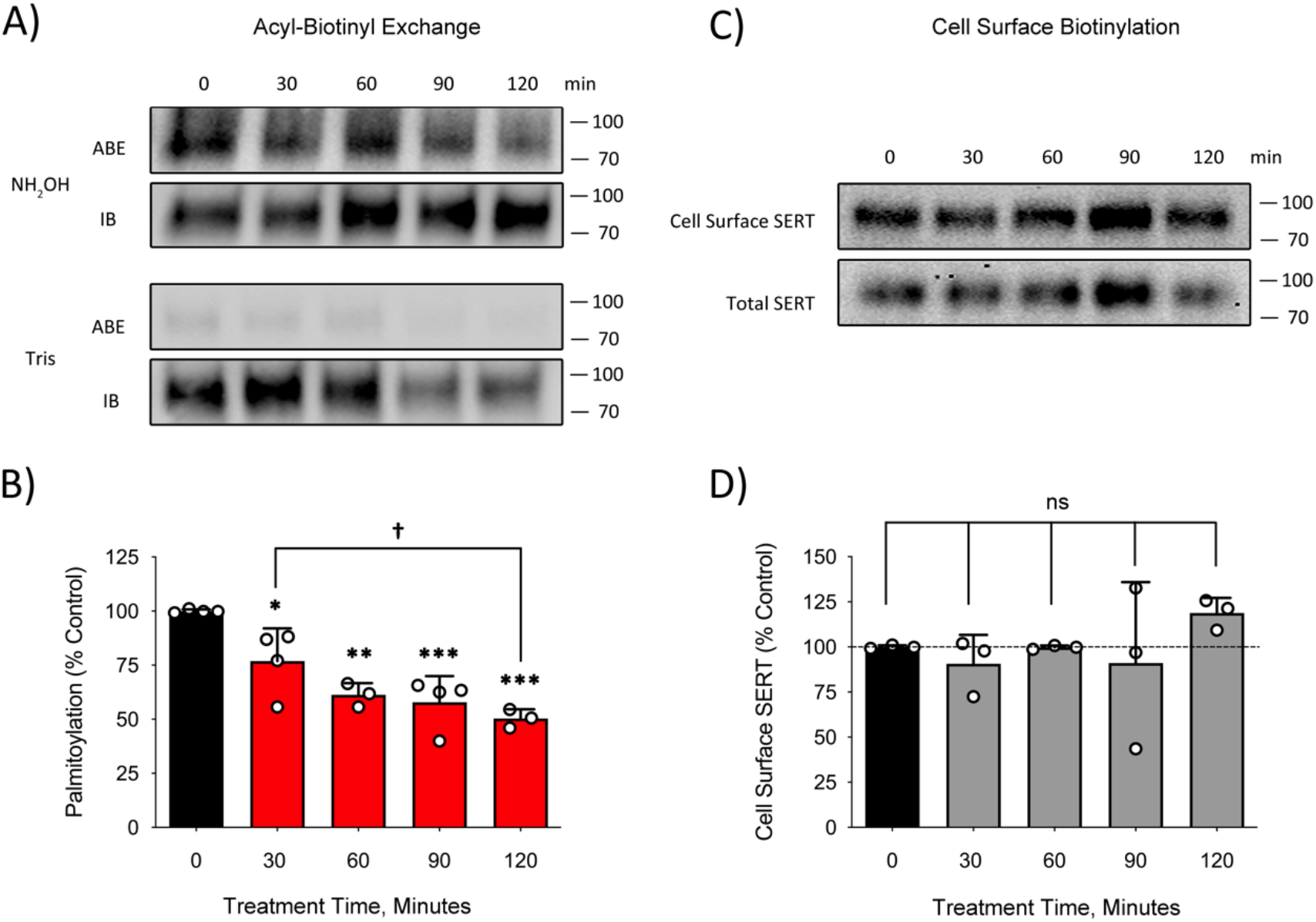
2BP inhibition of SERT palmitoylation does not change cell surface expression. *A,* HA-hSERT LLC-PK_1_ cells were treated with vehicle or 7.5 µM 2-BP for the indicted times and underwent ABE followed by SDS-PAGE and Immunoblotting for determining palmitoylation levels. M_r_ markers for all blots are shown at right. *B*, Quantification of palmitoylation intensity normalized to Total SERT levels (mean ± SD of three or more (n=3-4) independent experiments expressed relative to 0-time control as 100%; * p < 0.05, ** p < 0.01, *** p < 0.001 versus 0-time control; ^†^ p < 0.05 versus 30 min treatment; one-way ANOVA with Tukey post hoc test). *C,* Cell surface biotinylation of cell treated with vehicle or 7.5 µM 2-BP for the indicted times followed by SDS-PAGE and immunoblotting for hSERT. D, Quantification of surface SERT intensity (mean ± SD of three independent experiments relative to control normalized to 100%.; ns, p >0.05, one-way ANOVA with Tukey post hoc test).

After we identified a suitable concentration and time point for 2BP inhibition of palmitoylation with no impact on cellular SERT protein levels, we determined if changes in palmitoylation are accompanied by changes in trafficking and/or activity. Under the same conditions as shown in Figure 2B, cells were challenged with 7.5 µM 2BP in identical 30-minute intervals followed by SERT cell surface biotinylation and assessment of total cellular SERT levels. In this analysis, there was no change in SERT surface levels (Fig. 2C) and no apparent change in total SERT expression through 120 min of 2BP treatment (Fig. 2D) (p > 0.05, One way ANOVA with Tukey post hoc test, n=3).

Although acute palmitoylation changes do not appear to control DAT (21) or SERT trafficking (Fig. 2), we have previously demonstrated that 2BP decreases DAT V_max_ independent of changes in DAT surface expression (21,23). Because we did not observe any changes in SERT trafficking when treated with 7.5 µM 2BP, we examined 5HT uptake in response to 2BP in cells expressing SERT for potential changes in kinetic activity that may be analogous to those observed for DAT. We conducted saturation analyses of SERT when treated for 30 min with 7.5 µM 2BP, the concentration where we found the first observable change in palmitoylation outlined by our ABE data (Fig. 2A). This study revealed that 30 min of 7.5 µM 2BP treatment reduced SERT V_max_ to 61.7 ± 3.1% relative to vehicle control when normalized to SERT surface levels determined in parallel (Fig. 3A and B) (p<0.0001 via Student’s t-test, n=3). Outlined in Table 1, SERT V_max_ in the presence of vehicle (control) was 27.2 ± 2.6 pmol/min/mg while 2BP treatment reduced V_max_ to 17.2 ± 1.1 pmol/min/mg (p<0.001 via Student’s t-test for independent samples, n=3). No change in the K_m_ for 5-HT was observed (*Table 1*). Similar to the functional consequences of DAT palmitoylation, these results demonstrate that acute inhibition of SERT palmitoylation with 2BP downregulates SERT transport kinetics.

**TABLE 1.**
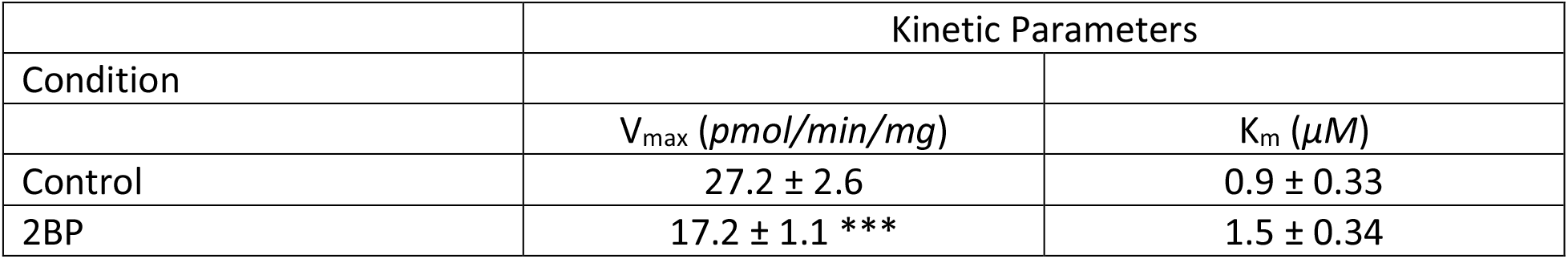
Kinetic parameters of hSERT after 2BP treatment. HA-hSERT LLC-PK_1_ cells treated with vehicle (control) or 7.5 µM 2BP for 30 minutes underwent 5HT transport saturation analysis (Fig. 3). Data are presented as mean ± SD for three independent experiments performed in triplicate. *** p<0.001 relative to control (Student’s t-test for independent samples).

**FIGURE 3.**
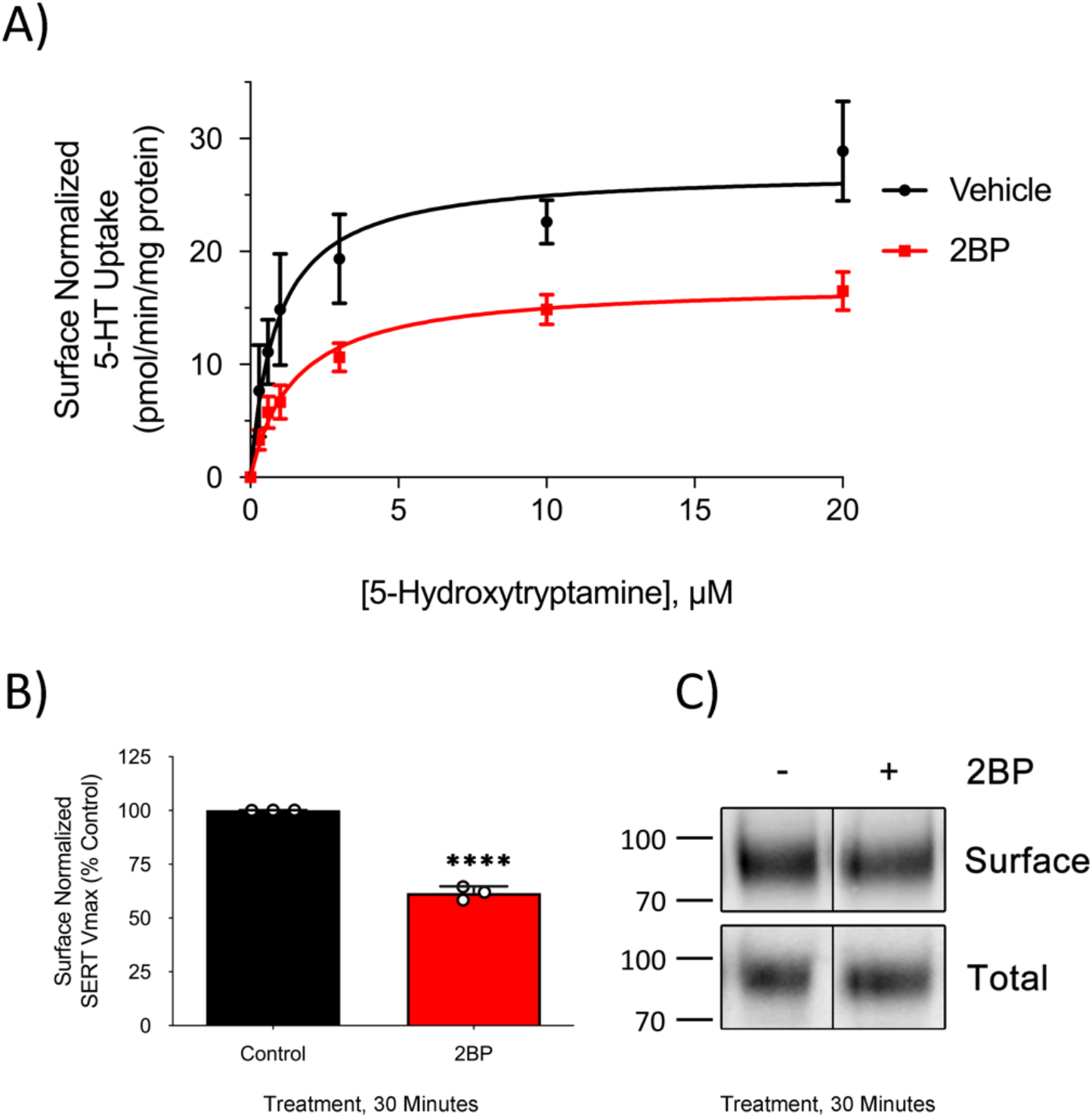
Inhibition of 5HT transport V_max_ by 2BP is independent of losses in cell surface SERT. HA-hSERT LLC-PK_1_ cells treated with vehicle or 7.5 µM 2BP for 30 minutes were subjected to 5-HT transport saturation analysis. *A*, Saturation analysis was performed and normalized to total cellular protein and SERT cell surface levels. *B,* Surface normalized SERT V_max_ (mean ± SD for three independent experiments performed in triplicate relative to control normalized to 100%). **** p<0.0001 relative to control (Student’s t-test for independent samples compared to control). *C*, Representative immunoblot of cell surface HA-hSERT and total SERT protein levels in 3 independent experiments after 30 minute treatments with vehicle or 7.5 µM 2BP. Black lines (|) within the blot image in panel C indicate removal of duplicate lanes. M_r_ markers are shown at the left of the blot in panel C.

Overall, when challenged with 2BP under acute conditions (7.5 µM 2BP for 30 min), reductions in palmitoylation did not alter the cell surface expression of SERT but did resulted in a decreased V_max_, that was independent of cell surface levels. This identifies palmitoylation as a major regulator in SERT function by directly changing the kinetic profile of the transporter, without altering the population of SERT at the cellular surface.

### Chronic Inhibition of SERT Palmitoylation Induces Loss of Total SERT Protein

We have demonstrated that chronic inhibition of DAT palmitoylation with 2BP, or site-directed mutagenesis of palmitoylated-cysteines, decreases total DAT protein and often produces DAT degradation fragments (21). To address if chronic inhibition of palmitoylation would impact SERT regulation and stability, we challenged LLC-PK_1_ cells expressing HA-hSERT with progressively increasing 2BP concentrations for 18 h. Because 2BP is a universal and irreversible inhibitor of the DHHC enzyme family, we hypothesized that a concentration-dependent reduction in palmitoylation would resemble our time-dependent findings in Figure 2. ABE analysis (Fig. 4A) revealed that palmitoylation of SERT was reduced with 10 µM 2BP to 71.6 ± 12.5% compared to vehicle control (p<0.001 via ANOVA with Tukey post-test, n=8) which continued with statistical significance through 50 µM 2BP (47.1 ± 21.8%, p<0.001 via ANOVA with Tukey post-test, n=3). Although we observed the anticipated concentration-dependent decrease in SERT palmitoylation, we were puzzled why SERT palmitoylation was not significantly lowered at concentrations lower than 10 µM 2BP with 18 h incubations. This was unexpected because our previous studies revealed that 7.5 µM 2BP inhibits SERT palmitoylation within 30 min (Fig. 2A).

**FIGURE 4.**
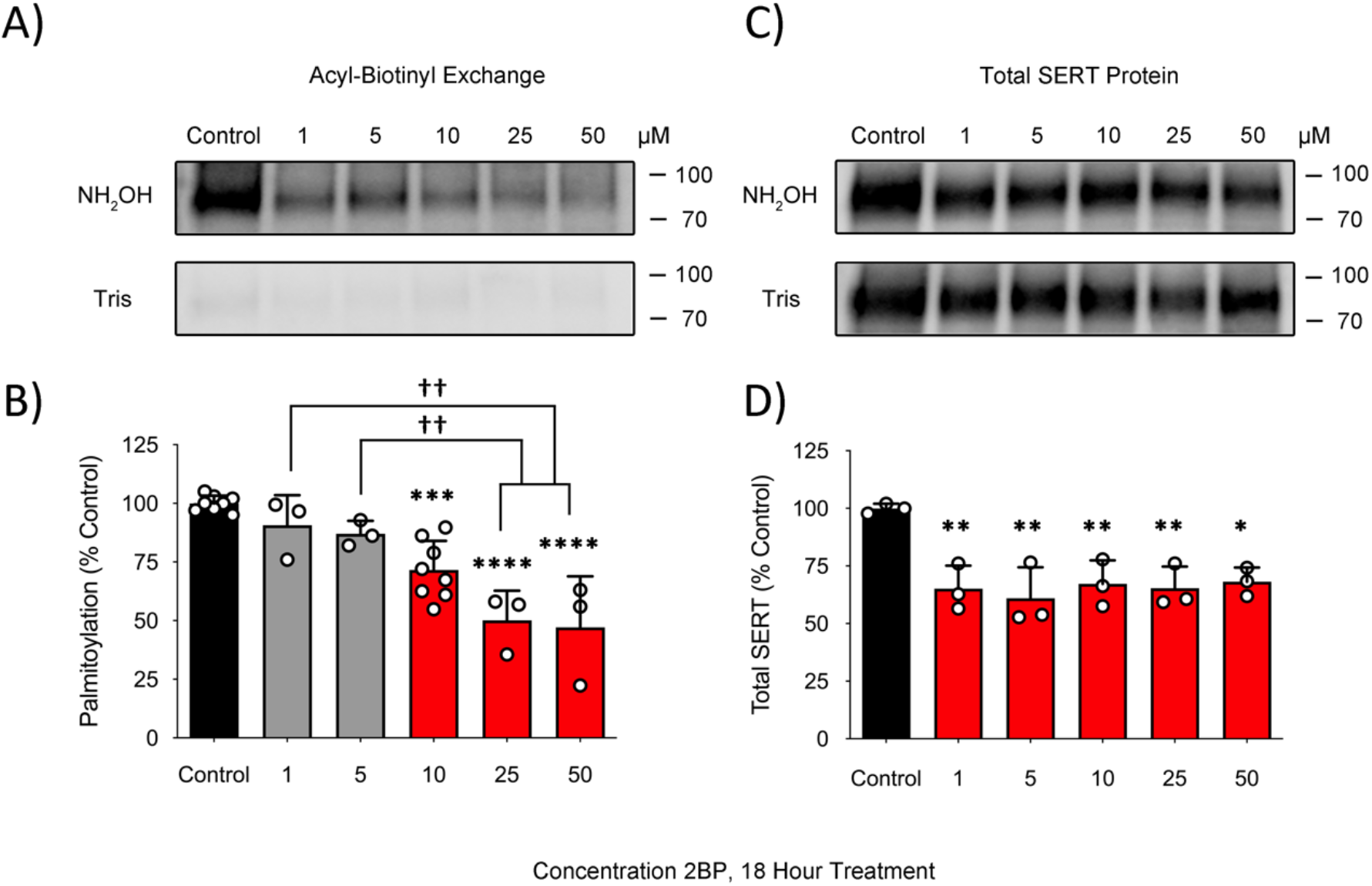
Chronic inhibition of SERT palmitoylation by 2BP is accompanied with a loss of cellular SERT protein. HA-hSERT LLC-PK_1_ cells were treated with the indicated concentrations of 2BP for 18 hours followed by assessment of hSERT palmitoylation levels by ABE and total hSERT levels by immunobloting. *A,* Representative immunoblot of hSERT following ABE analysis from three or more independent experiments. M_r_ markers for all blots are shown at the right. *B*, Quantification of palmitoylation normalized to total SERT present after each treatment (mean ± SD of three or more independent experiments (n=3-8) performed relative to control normalized to 100%.) **** p < 0.0001, *** p < 0.001, versus Control, ^††^ p < 0.01 versus the indicated concentrations (one-way ANOVA with Tukey post hoc test). *C,* Representative immunoblot of total HA-hSERT present with vehicle or 2BP treatments from three independent experiments. D, Quantification of total HA-hSERT protein in C (mean ± SD of three independent experiments relative to control normalized to 100%.) ** p < 0.01 versus control, * p < 0.05 versus control (one-way ANOVA with Tukey post hoc test).

Futher examination revealed that 2BP concentrations as low as 1 µM for 18 h induced a striking reduction in total SERT levels to 65.1 ± 9.9% of control which remained at 68.2 ± 6.2% of baseline through 50 µM of 2BP treatment when compared to vehicle control (Fig. 4C and 4D) (p<0.01, p<0.05 via ANOVA with Tukey post hoc test, respectively). In the ABE analysis, palmitoylated SERT quantification in NH_2_OH-treated fractions are normalized to total SERT protein present in the sample applied to the NeutrAvidin resin to account for potential losses of protein during the assay or treatment condition. Because total SERT levels are reduced after 18 h treatments even with lower 2BP concentrations (Fig. 4C and 4D), this resulted in quantified palmitoylation levels that were not different between vehicle control at 1 and 5 µM 2BP treatment conditions. However, it is likely that inhibition and subsequent reduction in SERT palmitoylation was the inciting mechanism of loss in total SERT protein similar to our findings previously reported for DAT (21).

### Escitalopram Decreases SERT Palmitoylation

We hypothesized that the reported reversible nature of SERT trafficking, when treated with escitalopram (14), was palmitoylation-dependent. In our acute time experiments we did not see a connection between changes in palmitoylation and trafficking properties of SERT (Fig. 2). However, palmitoylation is well known to regulate trafficking dependent properties of numerous proteins, altering compartment and cell surface localization. Because we also saw a loss in total SERT levels following 18 h of 2BP treatment, we hypothesized that SERTs state-of-palmitoylation may play a role in SERT processing and trafficking.

To test if escitalopram would affect SERT palmitoylation, we treated LLC-PK_1_ cells expressing HA-hSERT with 500 nM escitalopram for 3 h and subjected membranes isolated from these cells to ABE analysis. The concentration of escitalopram was derived from literature reports on maximal effects of escitalopram on SERT transport parameters in HEK293 cells via [^3^H]escitalopram binding and [^3^H]5HT saturation analyses (24). Likewise, 500 nM is within the range of steady-state plasma levels (95-720 nM) in individuals being treated with escitalopram 30-60 mg/day (14,25). Cells were treated with either 500 nM escitalopram or 7.5 µM 2BP as a positive control and were subjected to ABE followed by SDS-PAGE and immunoblotting for SERT. In four independent experiments, we found that SERT palmitoylation was reduced to 67.9 ± 14.4% of control by 2BP and 65.3 ± 6.2% of control by escitalopram (Fig. 5A and 5B) (p<0.00001 via ANOVA with Tukey post-test, n=4). Quantification of palmitoylation levels (Fig. 5A, *ABE*) were normalized to total SERT protein levels present (Fig 5A, *IB*) which accounted for any potential loss of protein in the ABE assay or during treatments. These results indicate that escitalopram reduces SERT palmitoylation to a level similar to that found with 2BP treatment.

**FIGURE 5.**
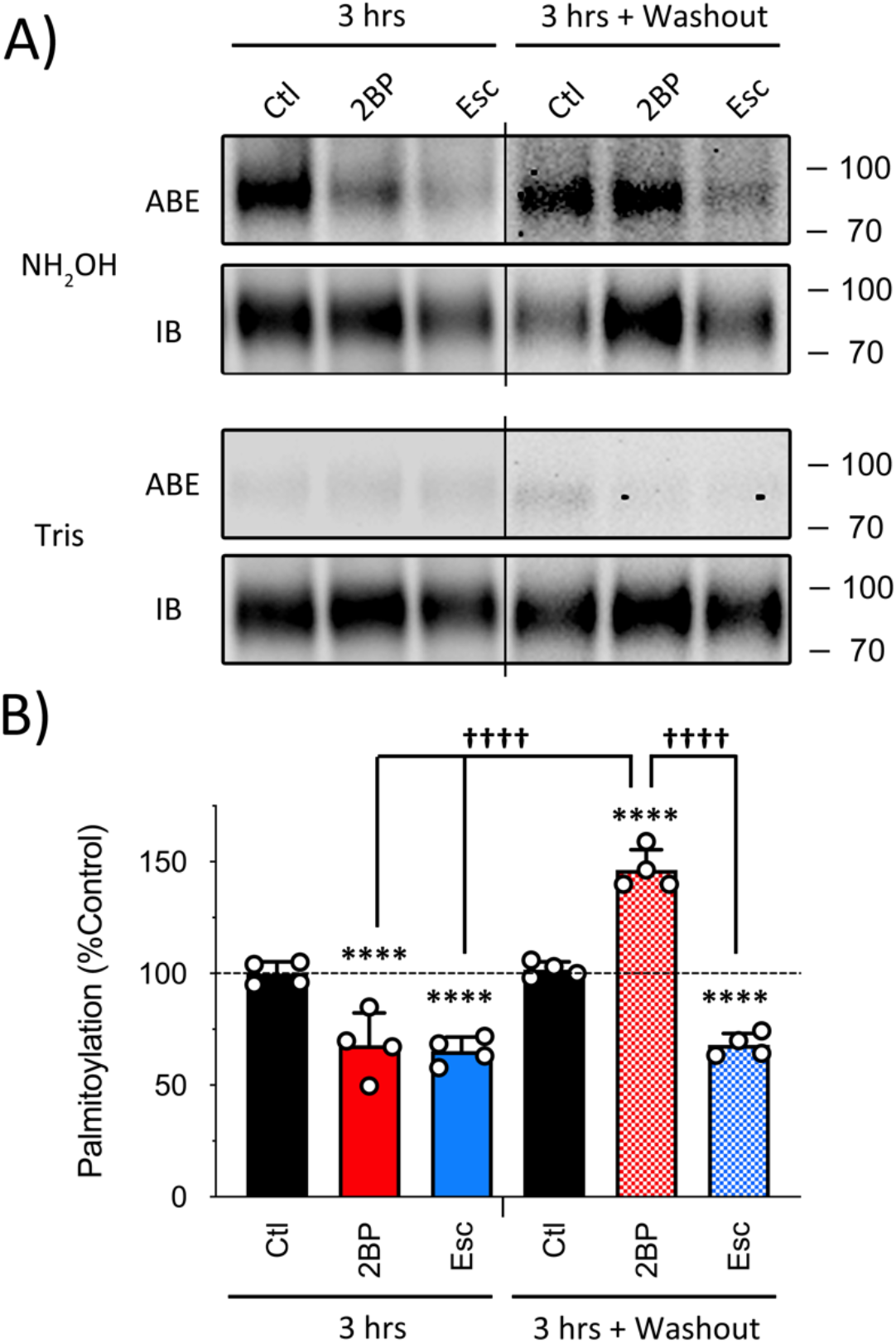
Escitalopram decreases palmitoylation of SERT. Cells were treated with vehicle, 7.5 µM 2BP or 500 nM S-citalopram for 3 h and were either immediately assessed for palmitoylation by ABE or washed twice and reincubated for an additional 3 h in drug-free media followed by assessment of palmitoylation levels using ABE. *A,* Representative immunoblot blot of four independent experiments (n=4) under the indicated conditions. Extended black lines (|) on top and bottom of images in panel A indicate images from separate blots. *B,* Quantification of palmitoylation (mean ± SD of four independent experiments relative to control (Ctl) normalized to 100%.) **** p < 0.0001 versus Ctl, ^††††^ p < 0.0001 between indicated treatment conditions (one-way ANOVA with Tukey post hoc test).

Our next step was to determine if this process could be reversed as described by Lau et. al. (14). To test this, we investigated palmitoylation of SERT in treatment conditions where the drugs were washed away using a protocol derived from Lau et. al. In these studies, cells were treated for 3 h with 500 nM escitalopram or 7.5 µM 2BP as a positive control, followed by two washes with drug-free buffer, then reincubation for an additional 3 h in drug-free buffer. These cells were then harvested, underwent membrane preparations, and ABE analysis. Following ABE, we identified that when cells were treated with 7.5 µM 2BP, and extensively washed, SERT palmitoylation was elevated to 146.3 ± 9% of baseline (Fig. 5, p<0.0001 via ANOVA with Tukey post-test, n=4). In comparison, after the wash and re-incubation cycle, SERT palmitoylation in escitalopram treated cells remained at 68 ± 5.2% of baseline (p<0.0001 via ANOVA with Tukey post-test, n=4), and unchanged from that observed in pre-wash conditions (Fig. 5). Unexpectedly, these results demonstrate that 2BP inhibition of SERT palmitoylation is sensitive to washout with a rebound in SERT palmitoylation levels. Likewise, these results also demonstrate that washout of extracellular escitalopram does not restore palmitoylation levels as hypothesized but maintains a long-term effect in LLC-PK_1_ cells.

### Escitalopram Decreases SERT Surface Expression and Transport Capacity

To investigate if 3h 2BP and escitalopram treatments that diminished palmitoylation was accompanied by decreased SERT surface expression and transport capacity, we performed cell surface biotinylation and 5HT uptake analysis on LLC-K_1_ cells expressing HA-hSERT treated under the same conditions used for the palmitoylation assessment shown in Figure 5. In this analysis we found that SERT surface expression was reduced to 65.4 ± 15.3% of control by 2BP (Fig. 6A and B, p<0.01 via ANOVA with Tukey post-test, n=6) and to 72.3 ± 13.2% by escitalopram after 3 h of treatment (*Fig. 6A and 6B,* p<0.05 via ANOVA with Tukey post-test, n=5). In washout conditions, cell surface SERT levels were not recovered after washout in either 2BP or escitalopram treatment conditions, where SERT surface levels remained decreased (2BP, 66.3 ± 18.8% of control; escitalopram, 51 ± 15% of control), under both treatment conditions (Fig. 6A and 6B, p<0.01 and p<0.001 respectively via Tukey post-test, n=3-4).

**FIGURE 6.**
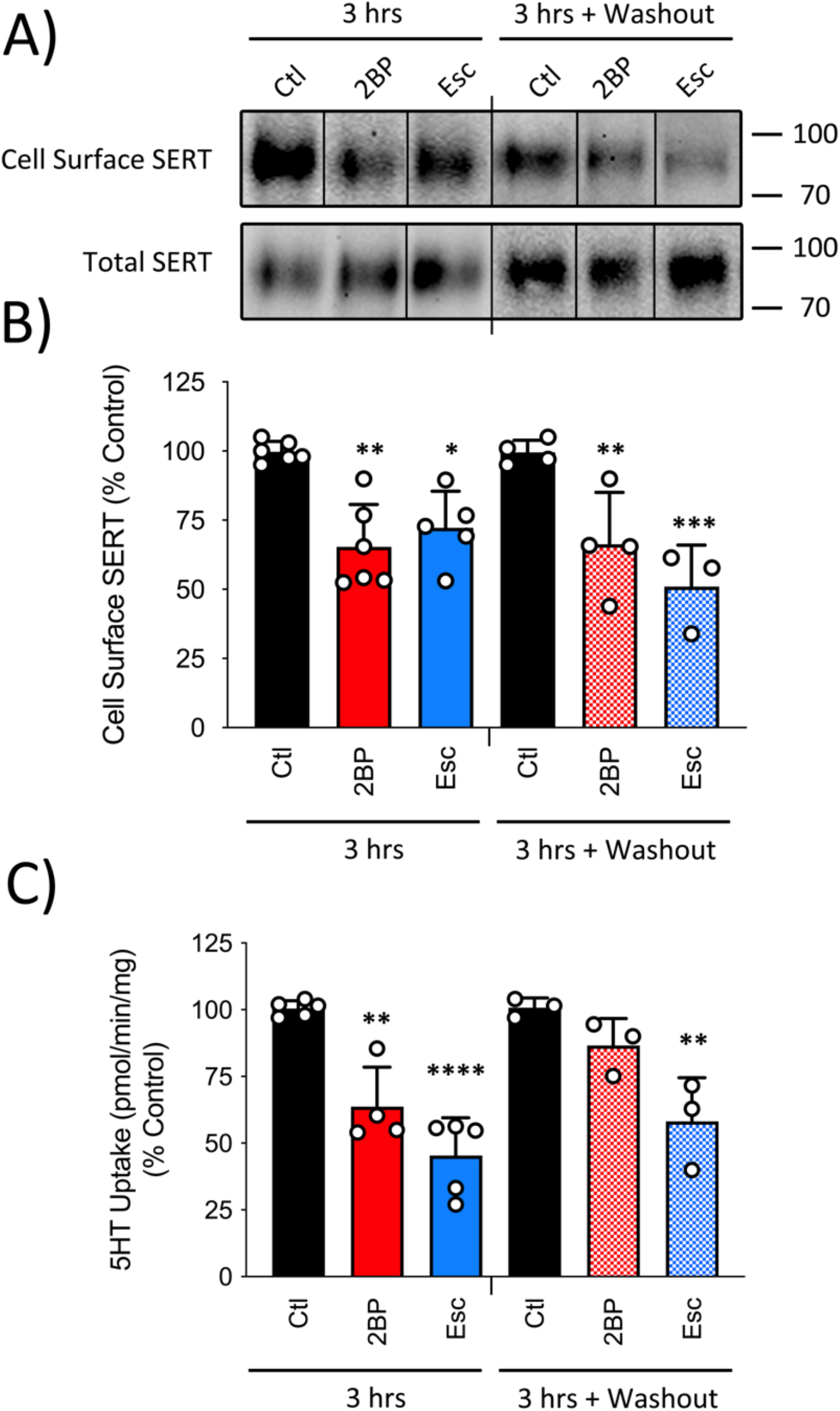
Escitalopram decreases SERT surface levels and transport capacity consistent with decreased palmitoylation. LLC-PK_1_ cells expressing HA-hSERT were treated with vehicle, 7.5 µM 2BP or 500 nM S-citalopram for 3 h followed by analysis for cell surface HA-hSERT expression *via* cell surface biotinylation (A and B) and 5HT transport capacity (C). Cells treated in parallel were washed twice and re-incubated for an additional 3 h in drug-free media followed by assessment of cell surface expression (A and B) and 5HT transport capacity (C). *A,* Representative immunoblot of cell surface and total HA-hSERT levels under the indicated conditions in four or more independent experiments (n=4-6). Extended black lines (|) on the top and bottom of panel A indicate two separate blots/experiments while black lines within the boundaries of the blot indicate the removal of duplicate lanes or rearrangement of lane images from the same blot. M_r_ markers for all blots are shown at right. *B,* Quantification of HA-hSERT cell surface immunoblots (mean ± SD of four or more independent experiments (n=4-6) relative to control (Ctl) normalized to 100%.) * p< 0.05, ** p <0.01, *** p<0.001 versus Ctl (One-way ANOVA with Tukey post hoc test). *C,* Quantification of HA-hSERT 5HT uptake (mean ± SD of three or more independent experiments (n=3-5) relative to control (Ctl) normalized to 100%.) * p< 0.05, ** p <0.01, *** p<0.001 versus Ctl (One-way ANOVA with Tukey post hoc test).

Following cell surface analysis, we performed 5HT uptake using these same treatment and washout conditions. Both 2BP and escitalopram treatments for 3 h resulted in decreased 5HT transport capacity (Fig 6C;. 63.7 ± 14.8% and 45.4 ± 14.1% of control, respectively) that was consistent with reduced SERT palmitoylation (Fig. 5) and surface levels (Fig. 6A and 6B).

In contrast, with 2BP treatment and a 3 h washout, SERT transport capacity rebounded to the control level (86.6 ± 10.1%; p>0.05 via ANOVA with Tukey post-test, n=3) while escitalopram treated cells following washout continued to display decreased 5HT transport (58.1 ± 16.4% of control (Fig.6C; p<0.01 via ANOVA with Tukey post-test, n=3). Escitalopram is a reversible competitive inhibitor of SERT and literature reports indicate it is readily removed under *in vitro* treatments following washout with recovery of 5HT transport capacity (26,27). To determine if the reduced transport capacity that persists after escitalopram washout was due to the presence of residual escitalopram and continued direct blockade of SERT, we performed a standard uptake assay with 10 µM 5HT after a brief treatment with 500 nM escitalopram or 10 µM cocaine as a control followed by washout just prior to uptake initiation. In both conditions, uptake after washout not reduced from controls (data not shown), suggesting that the majority of reduction in 5HT uptake demonstrated in Figure 6C was due to persistent palmitoylation and cell surface expression effects, and not a direct consequence of residual direct blockade by escitalopram. Collectively, these data generate many questions into the regulation of SERT via palmitoylation under differential therapeutic conditions. However, our results suggest that palmitoylation modulates trafficking and cell surface localization of SERT and is sensitive to inhibition by the DHHC inhibitor 2BP and the SSRI escitalopram.

## Discussion

Over the last 30 years, a large volume of evidence has characterized the regulation of monoamine transporters by post-translational modifications. These studies have revealed a means for cellular signaling to dynamically regulate synaptic communication and allow near-immediate physiologic responsiveness. Ultimately, synaptic homeostasis is maintained by these mechanisms and when they are dysregulated can promote the pathogenesis of disease.

SERT function is regulated by numerous signaling pathways that serve to maintain 5HT homeostasis in accordance with physiological demands. Evidence suggests that SERT is modified *via* post-translational modifications like phosphorylation (1,3,28), glycosylation (4,5), and ubiquitylation (29,30) that serve to coordinate SERT kinetic activity, localization, surface expression, and degradation in platelets, endothelial cells, and serotonergic neurons (1-3,6,28,31,32). Previous studies have reported that activation of PKC with β-PMA induces a time-dependent effect on SERT function that occurs in two phases. The first phase occurs at 5 min of PKC activation decreasing 5HT transport capacity without decreasing surface SERT levels, while the second phase occurs 30 min after activation, promoting the internalization of surface SERT (1). DAT and NET appear to share a similar method of regulation, with activation of PKC driving phosphorylation of both proteins and acutely decreasing their kinetic activity and surface expression (1,33-36). With DAT, our lab has demonstrated that palmitoylation levels are reciprocal of phosphorylation levels, with an enhanced palmitoylation state increasing kinetic activity and total DAT protein while driving phosphorylation by PKC activation decreases these processes (37). This increasing pool of evidence that monoamine transporters are regulated by post-translational modifications directed our attention to investigating if SERT is also palmitoylated like its monoamine cousin DAT.

Although our results demonstrated SERT is a palmitoylated protein, the modes of regulation mediated by palmitoylation were unclear and could be similar and/or different from DAT based on the location of the modified cysteines, the specific DHHC enzymes involved and their subcellular location. Several intracellular cysteines in the peptide chains of SERT and DAT, which serve as potential sites for S-palmitoylation, share a general sequence homology especially in the intracellular loops. We currently understand that cysteine 580 (C580) found at the DAT cytoplasmic / transmembrane domain 12 interface is palmitoylated (21), and a second palmitoylation site remains unidentified but is hypothesized to be Cys6 in the DAT N-terminus (38,39). In this respect, DAT and SERT both contain cytoplasmically accessible cysteine residues in relatively close proximity on their N- and C-termini. For the SERT N- and C-termini, cysteines are found at residues C15 and C22 in the N-terminus and residue C622 of the C-terminus which was identified in a pamitoylated peptide from a site specific analysis of neuronal protein S-acylation (40). The similarity in organization of these intracellular cysteines between SERT and DAT suggested to us that SERT regulation may be similar to DAT. Our findings demonstrate that there are acute and long-term distinctions in which palmitoylation regulates SERT, consistent with a previously reported biphasic effect mechanism (1), with short-term intervals using lower concentrations of 2BP being different than chronic conditions using higher concentrations of 2BP or longer incubation periods.

As demonstrated in our working model (*Fig. 7*), we found that acute 2BP treatment led to a strong reduction in 5HT transport V_max_ with no change in K_m_, surface, and total SERT expression (Fig. 7A and 7B. This suggests that, in acute timeframes, palmitoylation regulates SERT transport kinetics without altering SERT surface expression or total transporter levels. Notably, our previous findings with DAT outline similar outcomes, with alterations in palmitoylation controlling DA transport V_max_ without changes in DAT surface expression or total DAT levels (21,23,37). This finding is consistent with our hypothesis that acute changes in palmitoylation have similar regulatory properties for SERT and DAT and suggests that SERT may be palmitoylated on C-terminal cysteine (C622) and one or more N-terminal cysteine residues (C15 and/or C22). However, this is currently unknown, as we have not yet examined individual cysteines of SERT for palmitoylation.

**FIGURE 7.**
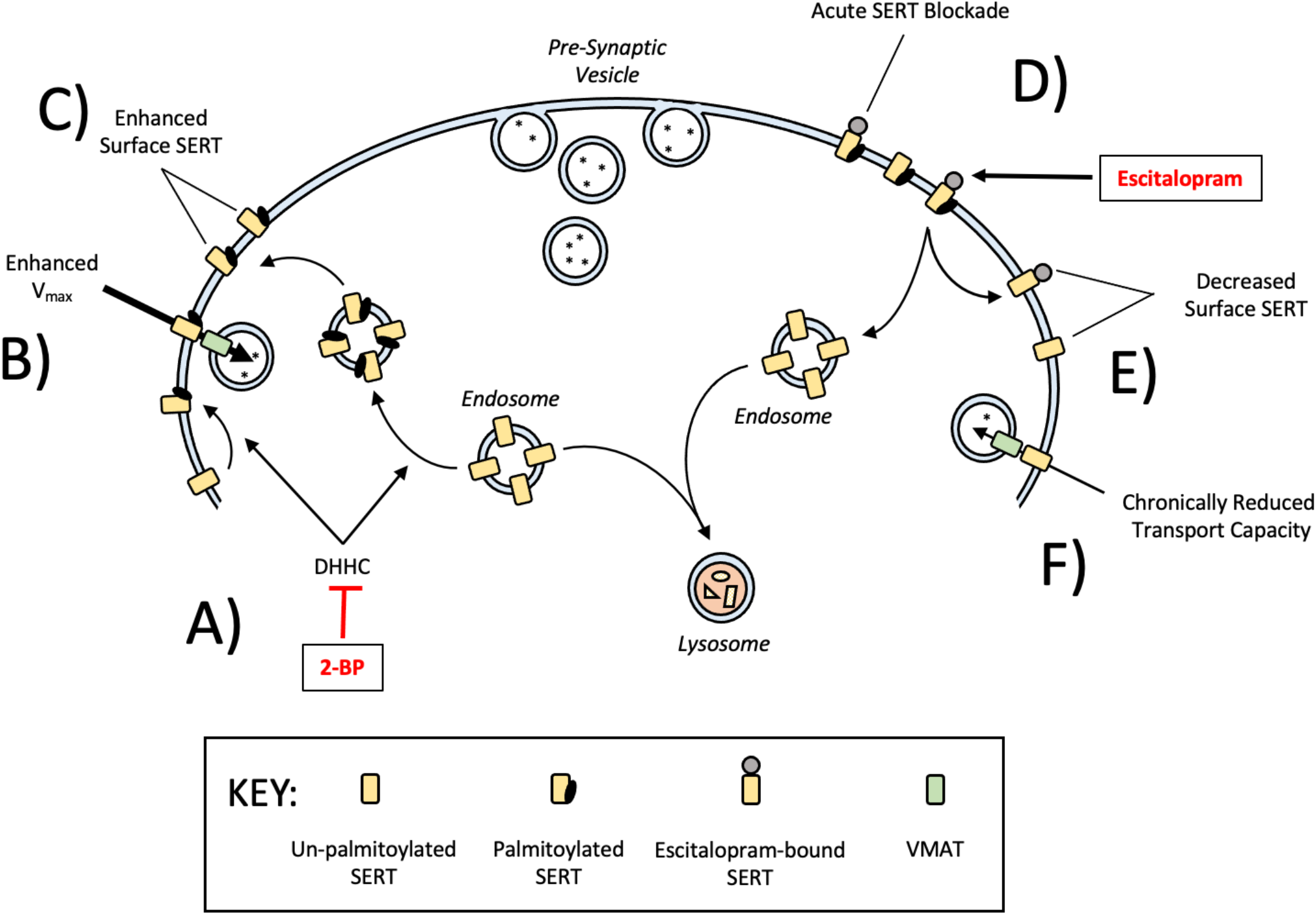
Model for regulation of SERT by palmitoylation. *A,* 2BP sensitive DHHC enzymes palmitoylate SERT, enhancing V_max_, cell surface recruitment, and oppose loss of total SERT. *B*, Acute palmitoylation enhances SERT V_max_ independent of cell surface levels. *C,* Palmitoylation also promotes the recruitment of SERT to the cellular surface in the long-term opposing internalization. *D,* Escitalopram decreases SERT palmitoylation levels that is not recoverable upon drug removal. It is currently unclear if escitalopram binds to SERT in its palmitoylated state and encourages removal of palmitoylation, if escitalopram binds to SERT in its un-palmitoylated state and prevents palmitoylation, or both. *E*, Following exposure to escitalopram or 2BP, SERT palmitoylation and surface levels are decreased by internalization, inhibition of surface recruitment, or both. *F,* SERT palmitoylation and surface expression through these mechanisms results in decreased 5HT transport capacity that persists after drug removal.

It seems likely that SERT may undergo palmitoylation in response to acute changes in physiologic stimuli. Acutely, SERT kinetic activity may be enhanced to modulate acute elevations in synaptic 5HT and prevent abnormally increased 5HT signaling and spillover. This makes sense, as SERT undergoes several other post-translational modifications that are responsive to physiologic stimuli; the most notable is PKC-driven phosphorylation that rapidly decreases SERT kinetic activity (1). Likewise, it has been determined that SERT is a target for p38 MAPK in response to p38 MAPK (41,42) and adenosine receptor activation (228). In acute timeframes, activation of the A3 adenosine receptor induces an increase in 5HT uptake V_max_ (41). These findings strongly suggest that the dynamic nature of palmitoylation may be responsive to other physiologic stimuli, functioning to acutely regulate extracellular 5HT concentrations following release. However, the DHHC machinery that catalyzes the attachment of palmitate to proteins have yet to be extensively examined under the influence of various physiologic stimuli but are very likely to be physiologically responsive. In this regard, the relatively rapid rebound of SERT palmitoylation after washout of DHHC inhibitor, 2BP, without recovery of surface levels suggests the potential for a rapid kinetic response by this mechanism.

When treated with higher concentrations of 2BP in longer time intervals, we observed losses of total SERT protein that were consistent with losses in palmitoylation. These data suggest that palmitoylation is responsible for maintaining long-term stability of SERT either by opposing degradation as seen with DAT (21,23) or by supporting biogenesis as seen with cystic fibrosis transmembrane conductance regulator (CFTR) protein (19). Previously, we demonstrated that total DAT levels are sensitive to inhibition with prolonged 2BP treatment that resemble our findings with SERT (Fig. 4B). Specifically, when DAT palmitoylation was reduced after treatment with high 2BP concentrations for longer time intervals, or by C580A mutation, we observed dramatic losses in total transporter expression with the production of degradation fragments (21). In the current study, we did not detect any SERT fragments, suggesting the possibility that palmitoylation may function co-translationally to enhance SERT biogenesis. In this mechanism, it is possible that inhibition of SERT palmitoylation at the co-translational level would result in decreased SERT protein expression, without evidence of lysosomal degradation (Fig 7C). This mechanism of long-term control seems likely due to the role palmitoylation plays in modulating the biogenesis of other proteins. In a viral model, the SARS-CoV-2 Spike (S) protein undergoes constitutive palmitoylation at the ER level and is required for infectivity (43,44). The S protein contains four cysteine-rich clusters, with clusters 1 and 2 serving as the primary sites for palmitoylation (43). When these sites are disrupted by mutation, ER-to-Golgi trafficking is suppressed. Likewise, S protein membrane fusion activity is dramatically reduced (43). These results highlight the role of palmitoylation in S protein maturation and processing and suggests that SERT may utilize a similar method of post-translational processing with inhibition of this process leading to a decrease in total SERT levels.

Our data also suggest that palmitoylation likely controls post-translational trafficking of SERT at the endosomal level. In this regard, we found that three h of 2BP-mediated inhibition led to decreased SERT surface expression. As depicted in our working model (Fig. 7), we propose that palmitoylation has a time-based mechanism in modulating SERT activity, with 30 min of inhibition decreasing SERT V_max_ independent of surface levels (Fig. 7B), and 3 h of inhibition causing decreased SERT surface expression (Fig. 7D and 7E). It is possible that the reduction in surface SERT may be multi-faceted, with inhibition of palmitoylation causing cell surface SERT to be internalized and/or by preventing the recruitment of SERT to the surface from endosomal stores. Collectively, it seems likely that both of these processes reflect an overall reduction of SERT at the cellular surface, but further exploration is required.

In our previous work, we have characterized some of the DHHC family members that enhance DAT palmitoylation and subsequent functional consequences like transport capacity, surface, and total DAT expression (23). SERT and DAT share similar sequence homology in their internal core structure but vary considerably in amino acid composition and location of cysteine residues on the C- and N-termini. This suggests that palmitoylation may play similar and/or different roles in controlling dopaminergic and serotongergic homeostasis. In this mechanism, it’s possible that palmitoylation of SERT on unique cysteine residues, by a different collection of DHHCs localized in different subcellular locations, may lead to differences in physiologic and sub-cellular regulatory patterns that are distinct between SERT, DAT, NET, and other membrane transporters. This seems likely, as signaling physiology is dictated by tissue and cellular specificity in order to conduct normal physiological responses between organ systems. Overall, it seems plausible that DHHCs may palmitoylate SERT on different cysteine residues in distinct subcellular membrane locations to execute distinct outcomes, but further exploration to demonstrate this complex relationship is necessary.

Other studies have demonstrated that prolonged inhibition of neuronal SERT by escitalopram induces transporter internalization and translocation to the cellular soma (14,15,45). Remarkably, the total density of cellular SERT appears to decrease as well, with several studies reporting that SERT total expression and mRNA levels are decreased upon chronic exposure to escitalopram in human platelets and rat raphe neurons (14,15,46-48). Importantly, our interest was stimulated upon discovering reports that these processes could be reversed when the drug was removed (14). Palmitoylation is dynamic and reversible, allowing numerous cycles of addition and removal of palmitate to a protein throughout its lifetime, which led us to speculate that escitalopram may modulate SERT palmitoylation. This directed our attention to the possibility that the therapeutic mechanism for escitalopram may include palmitoylation-mediated internalization of SERT in addition to chronic elevations in 5HT and post synaptic signaling. This molecular explanation is supported by evidence of the lasting effects of antidepressant exposure on neonatal rodent behavior and serotonin circuitry (49) where the authors demonstrated SSRI induced delayed normal maturation of the serotonergic system that was a consequence of their SERT specific effects (49). Likewise, the effects of SSRI use during human pregnancy and the subsequent impact on neonatal outcomes has long been debated. In a case report with six neonates, SSRI withdrawal was evident between 2-3 weeks post-partum, a period in which the bioavailability of SSRIs are long expected to be decreased (41,50). In this post-exposure period, neonates demonstrated varying degrees of myoclonus, irritability, jitteriness, feeding problems, and others that are classically associated with serotonin syndrome (51). The evidence collected at the molecular and clinical levels supports the idea that SSRIs function to decrease SERT activity, expression and overall long-term serotonergic tone. Based on our findings, it seems likely that palmitoylation may be an important component in controlling the density and duration of SERT protein at the cellular surface, which subsequently seems important in dictating overall brain structure and neuronal networks associated with autism, neonatal SSRI withdrawal, and depression.

As with peripheral tissue, it is known that the brain has a complex interplay of extracellular matrices (ECM) that can be remodeled upon varying physiologic requirements. Studies suggest that changes in the composition of human, rodent, and zebrafish CNS ECM are associated with alterations in neuronal connectivity and may contribute to the cognitive and emotional attributes of depression (52). In rodents, a causal link has been identified between increased hippocampal ECM density and the cognitive deficits associated with chronic depression, suggesting that abnormal changes in the ECM may play a role in the development of major depression (53). In human platelets, it has been discovered that integrin αIIbβ3 interacts with the C-terminus of SERT, enhancing SERT surface expression (54). Integrins are well known to be the principal receptor used in linking the extracellular matrix and intracellular actin cytoskeleton, with disruption of this process resulting in numerous pathologies best understood in cancer development and defects in cellular migration like leukocyte adhesion deficiency (55,56). Although unexplored, it seems possible that escitalopram may function to alter neuronal connectivity through SERT-dependent extracellular remodeling with decreased SERT expression in the neuronal plasma membrane. This seems possible, as SSRIs have been reported to require at least one week to three months of constant dosing to achieve clinically observable neuro-cognitive effects, suggesting that the therapeutic potential for SSRIs require longer-term adaptations to the serotonergic system (11). This unique explanation provides a sufficient timeframe for ECM remodeling to coincide with clinically significant antidepressant effects.

Ultimately, our findings provide the first report of SERT as a palmitoyl-protein (57,58), and contribute a novel component to an already extensive pool of existing literature on post-translational modifications of monoamine transporters. The regulatory nature of post-translational modifications, including palmitoylation, are critical in maintaining physiologic homeostasis in response to cellular demands and environmental challenges. Many DHHC and APT enzymes are expressed in neuronal tissue and dynamically regulate the palmitoylation level of numerous proteins involved in neurotransmission and synaptic homeostasis (59-64). Importantly, escitalopram, a widely prescribed antidepressant, was found to decrease SERT palmitoylation, surface expression, and long-term transport capacity following washout. This suggests that escitalopram promotes an unpalmitoylated SERT state that may contribute to the clinical findings associated with SSRI withdrawal and the length of time required for SSRI efficacy. Our results suggest that regulation of SERT by palmitoylation is essential in maintaining serotonergic homeostasis *via* acute and chronic mechanisms and that disruption of this process either by disease or therapeutics may lead to clinically observable neurocognitive changes.

## Experimental procedures

### Cell Culture

Lilly Laboratory Porcine Kidney (LLC-PK_1_) cells stably expressing hemagglutinin (HA) tagged WT hSERT, and Human Embryonic Kidney-293 (HEK293) cells stably expressing WT hSERT were grown in Dulbecco’s Modified Eagle Medium (DMEM) containing 5% fetal bovine serum, 100 μg/mL penicillin/streptomycin, and supplemented with 400 μg/mL of Geneticin (G418) for maintenance of stable expression. Cells were maintained in a humidified incubator gassed with 5% CO_2_ at 37°C. SERT expression level was verified by sodium dodecyl sulfate-polyacrylamide gel electrophoresis (SDS-PAGE) and immunoblotting of the cellular lysates against Anti-HA (Biolegends) and anti-human (h) SERT (Mabtechnologies-ST51-2) specific antibodies.

### Membrane Preparation

LLC-PK_1_ cells expressing the indicated SERTs were grown in 100 mm plates to 85% confluency. Cells were washed twice with 3 mL of ice-cold Buffer B (0.25 mM sucrose, 10 mM triethanolamine, pH 7.8), scraped, and collected in 500 µL of Buffer B containing a protease inhibitor cocktail of 1 μM phenylmethylsulphonyl fluoride (PMSF) and 5 μM Ethylenediaminetetraacetic acid (EDTA) at 4°C and transferred to a 2 mL microcentrifuge tube. Cells were then pelleted *via* centrifugation at 3,000 *x g* for 5 min at 4^°^C, the supernatant fraction was removed, and the cell pellet was suspended in 1 mL of ice-cold Buffer C (0.25 M sucrose, 10 mM triethanolamine, 1 mM EDTA, 1 µM PMSF, pH 7.8) and subsequently homogenized *via* 30 strokes of the pestle in a Dounce homogenizer. Homogenates were cleared of cellular debris and nuclei by centrifugation at 800 *x g* for 10 min. The post-nuclear supernatant fraction was collected and centrifuged at 18,000 *x g* for 12 min at 4°C to pellet cell membranes. The resulting membrane pellet was suspended in 1mL of sucrose phosphate (SP) buffer (10 mM sodium phosphate, 0.32 M sucrose, pH 7.4 with 1 μM PMSF and 5 μM EDTA) and assayed for protein concentration.

### Acyl-Biotinyl Exchange (ABE)

The ABE method used as previously described was adapted from Wan et. al (22) where palmitoylated proteins are detected in three steps: (I) Free cysteine thiols are blocked; (II) thioester linked palmitoyl groups are removed by hydroxylamine (NH_2_OH); (III) the formerly palmitoylated and now newly generated sulfhydryl groups are biotinylated. Membranes prepared from LLC-PK_1_ cells expressing HA tagged-WT hSERT or HEK293 cells expressing WT hSERT were solubilized in 250 μL of lysis buffer (50 mM HEPES pH 7.0, 2% SDS (w/v), 1 mM EDTA) containing 25 mM N-ethylmaleimide (NEM), incubated for 20 minutes in a 37°C water bath and mixed end-over-end for at least 1 hour at ambient temperature. Proteins were precipitated *via* the addition of 1 mL acetone and centrifugation at 18,000 *x g* for 10 min. The protein pellet was resuspended in 250 µl lysis buffer containing 25 mM NEM and incubated for 1 hour at ambient temperature. This process was repeated a final time with incubation overnight with end-over-end mixing. NEM was removed *via* acetone precipitation/centrifugation and the protein pellet was resuspended in 250 μL 4SB buffer (50 mM Tris, 5 mM EDTA, 4% SDS, pH 7.4) and remnant NEM was removed by an additional acetone precipitation and centrifugation. The pellet was resuspended in 200 μL 4SB buffer and thioesterified palmitate molecules were removed by incubation with hydroxylamine (NH_2_OH); The sample was split into two equal aliquots (100 μL) with one diluted with 800 µl Tris-HCl, pH 8.0 (negative control) and the other diluted with 800 µL NH_2_OH (0.7 M final concentration) and incubated at ambient temperature for 30 min with end-over-end mixing. Both samples were treated with 100 μL of a sulfhydryl-specific biotinylating reagent, HPDP-biotin (0.4 mM final concentration) and incubated at ambient temperature with end-over-end mixing for 1 h. NH_2_OH and biotin reagents were removed by acetone protein precipitation with centrifugation at 18,000 *x g* and supernatant fraction aspiration. Protein pellets were then resuspended and solubilized in 150 μL 4SB, acetone precipitated with 600 µL, centrifuged at 18,000 *x g* followed with supernatant aspiration. The final pellet was suspended in 75 μL ABE lysis buffer. A 10 μL aliquot was set aside for determination of total SERT content by immunoblotting while 65 μL was diluted in 1500 μL Tris buffer and incubated with 50 μL of a 50% slurry of Neutravidin resin to affinity purify the biotinylated proteins/peptides overnight at 4°C with end-over-end mixing. Unbound proteins/peptides were washed away by three cycles of 8,000 *x g* centrifugation, removal of the supernatant fraction, and resuspension in 750 μL radioimmunoprecipitation assay buffer (RIPA: 1% Tx-100, 1% sodium deoxycholate, 0.1% SDS, 125 mM sodium phosphate, 150 mM NaCl, 2 mM EDTA, 50 mM NaF). Proteins and peptides were eluted from the final pellet by incubation in 2x Laemmli sample buffer (SB: 125 mM Tris-HCl, 20% glycerol, 4% SDS, 200 mM DTT, 0.005% bromophenol blue) for 20 min at ambient temperature. Samples were then subjected to SDS-PAGE and immunoblotted with mouse anti-HA primary antibody (Anti-HA) or mouse anti-human SERT primary antibody (ST51-2).

### Cell Surface Biotinylation

LLC-PK_1_ cells stably expressing HA-tagged hSERT were grown in 24-well plates until 80% confluent. After treatments, the cells were washed three times with ice-cold Hank’s balanced salt solution containing Mg^2+^ and Ca^2+^ (HBSS Mg-Ca: 1 mM MgSO_4_, 0.1 mM CaCl_2_, pH 7.4), and subsequently incubated twice with 0.5 mg/mL of membrane-impermeable sulfo-NHS-SS-biotin for 25 min on ice with rocking. The biotinylation reagent was removed by aspiration and the reaction was quenched by two sequential incubations with 100 mM glycine in HBSS Mg-Ca for 20 min on ice with rocking. Cells were washed with HBSS Mg-Ca and then lysed with 250 μL per well RIPA buffer containing protease inhibitor. Lysates from 4 identically treated wells were pooled and assayed for protein content. Equal protein from each lysate (100 μg in 750 µl of RIPA) was incubated with 50 µL of a 50% slurry of Neutravidin resin overnight at 4°C with end-over-end mixing. The protein bound resin was washed three times with RIPA buffer, and the bound protein was eluted with 32 μL of Laemmli SB followed by SDS-PAGE and immunoblotting for HA-SERT with Anti-HA antibody.

### Saturation Analysis

LLC-PK_1_ cells stably expressing HA-tagged hSERT were grown in 24-well plates until 80% confluent. The cells were washed with 0.5 mL 37°C Krebs Ringers HEPES (KRH: 120 mM NaCl, 5 mM KCl, 2 mM CaCl_2_, 1 mM MgCl_2_, 25 mM NaHCO_3_, 5.5 mM HEPES, 1 mM D-Glucose) buffer and re-incubated in KRH buffer with or without 7.5 μM 2BP for 30 min at 37^0^C. Immediately following 2BP incubation, 5HT uptake was conducted for 8 min at 37°C with 0.32, 0.6, 1, 3, 10, and 20 μM total 5HT containing 20 nM [^3^H]5HT; nonspecific uptake was determined in the presence of 100 µM s-citalopram. Cells were rapidly washed twice with 500 µL ice-cold KRH buffer and lysed with 1% Tx-100 for at least 20 min with rocking at ambient temperature. Radioactivity contained in lysates was assessed by liquid scintillation counting. Kinetic values were determined using Prism software, and V_max_ values were normalized to total cellular protein (pmol/min/mg) and transporter surface levels determined by surface biotinylation assays performed in parallel for each experiment.

### [^3^H]5HT Uptake Assay

LLC-PK_1_ cells stably expressing HA-tagged hSERT were grown in 24-well plates until 80% confluent and were treated with the indicated concentrations of 2BP or escitalopram prepared in DMSO or KRH respectively for 3 hours at 37°C followed by [^3^H]5HT uptake assay in KRH. Equal DMSO concentrations were used in appropriate vehicle controls and final DMSO concentrations per well were 1% or less, which by itself did not affect 5HT transport activity. Uptake assays were performed in triplicate for 8 min at 37°C with 3 µM 5HT (20 nM [^3^H]5HT) and nonspecific uptake was determined in the presence of 100 nM (-)escitalopram. Following uptake, the cells were rapidly washed three times with ice-cold KRH. Cells were then solubilized in 1% Triton X-100, and radioactivity contained in lysates was assessed by liquid scintillation counting.

### SDS-PAGE and Western Blotting

Proteins were denatured in Lammeli SB and were electrophoretically resolved using 4-20% polyacrylamide gels alongside a molecular weight protein standard. Proteins were then transferred onto polyvinylidene fluoride (PVDF) membrane and immunoblotted for SERT using anti-HA or anti-human SERT (ST51-2) antibodies diluted 1:1,000 in blocking buffer (3% bovine serum albumin, phosphate buffered saline (PBS: 0.137 M NaCl, 0.0027 M KCl, 0.01 M NaH_2_PO_4_, 0.0018 M KH_2_PO_4_) and incubated for at least two hours at ambient temperature or overnight at 4°C. After five washes for 5 min each with wash buffer (0.1% Tween 20, 1x PBS), the membrane was incubated for 45 minutes in alkaline phosphatase (AP)-linked anti-mouse IgG secondary antibody diluted 1:5,000 in blocking buffer followed by 5 additional 5 min washes. Protein bands were visualized by chemiluminescence using Immun-Star(tm) AP substrate (Bio-Rad) applied to the membrane and incubation for 5 min at ambient temperature. Band intensities were quantified using Quantity One® software (Bio-Rad).

## Data availability

The original contributions presented in the study are included in the article. Further inquiries can be directed to the corresponding author.

## Acknowledgements

This work was supported by NIH grants 2R15DA031991-02A1 and P20 GM103442 (IDeA) from INBRE of the NIGMS, NSF REU site grant 1852459, and the UND SMHS.

## Abreviations

The abbreviations used are:

LLC-PK_1_: Lilly lab porcine kidney cells
SERT: serotonin transporter
DAT: dopamine transporter
NET: norepinephrine transporter
CFTR: cystic fibrosis transmembrane conductance regulator
ABE: acyl biotinyl exchange
DHHC: palmitoyl acyltransferase
APT: Acyl protein thioesterase
5HT: 5-hydroxytryptamine
2BP: 2-bromopalmitate
PKC: protein kinase C
PMA: phorbol 12-myristate, 13-acetate
CNS: central nervous system
SSRI: serotonin selective reuptake inhibitor
MAO: monoamine oxidase
TCA: tricyclic antidepressant
MAPK: mitogen-activated protein kinases

